## Supplementary information for "ATP burst is the dominant driver of antibiotic lethality in *Mycobacterium smegmatis*"

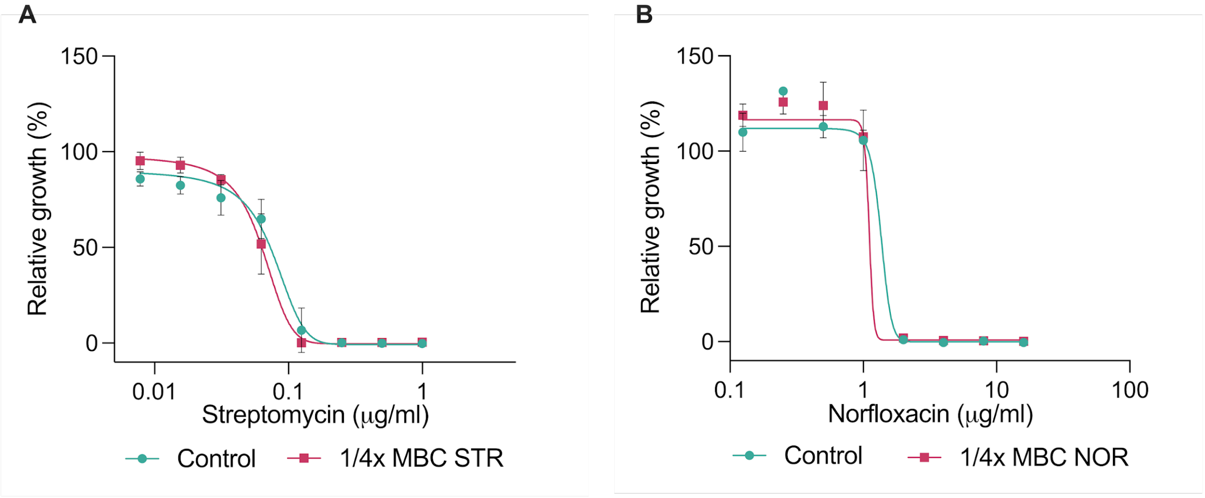


**S1.** Graphs depict the MIC of *M. smegmatis* grown in the presence and absence of ¼x $MBC_{99}$ of streptomycin (A) or norfloxacin (B) for 25 hours. Cells were harvested at 25 hour (recovery phase) and MIC was compared.

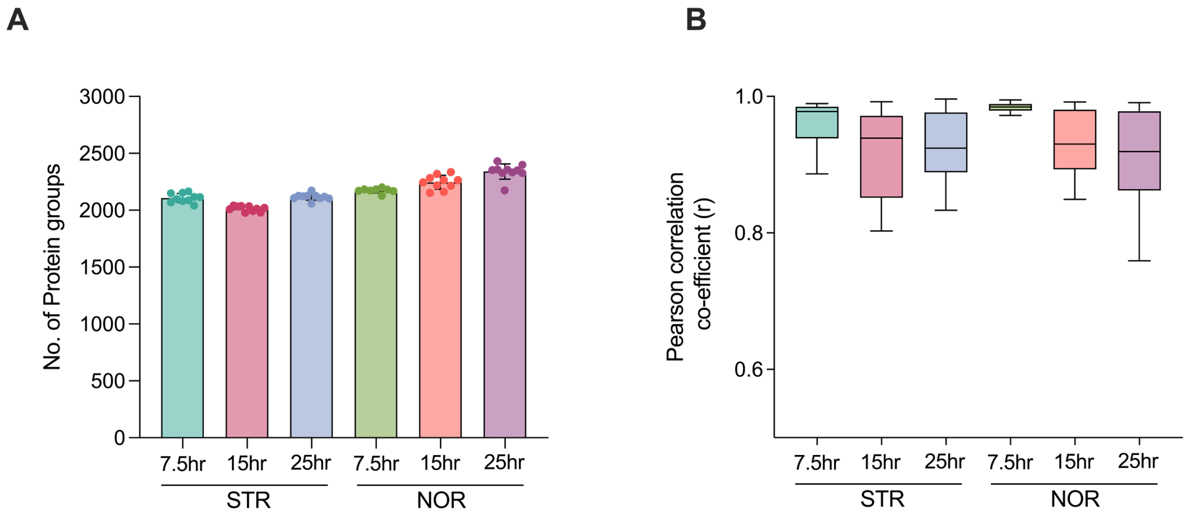


**S2.** Bar graph (A) shows the number of proteins identified from norfloxacin and streptomycin treatment at 7.5, 15, and 25 hours. Each dot represents the number of proteins identified per sample. Box plot (B) depicts the Pearson correlation analysis of the LFQ intensities among all 10 replicates of each set.

A

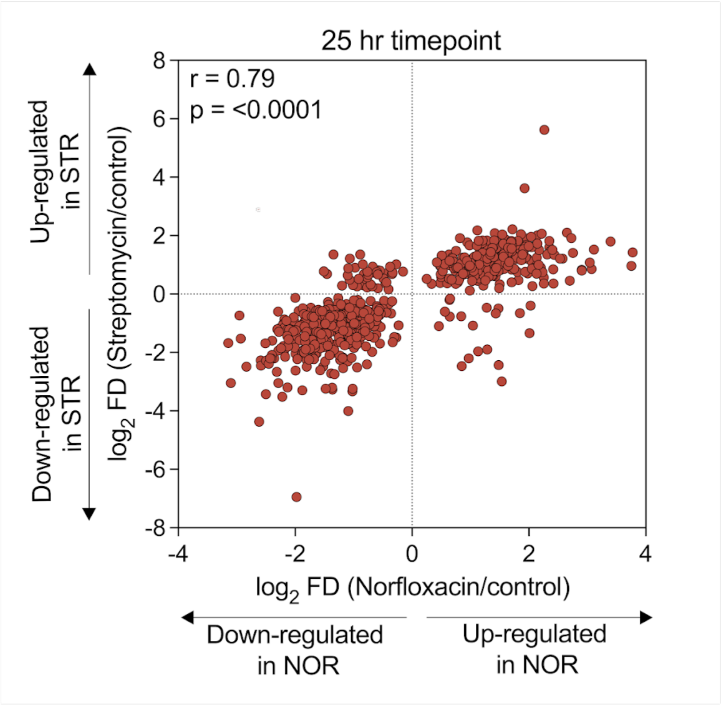


B

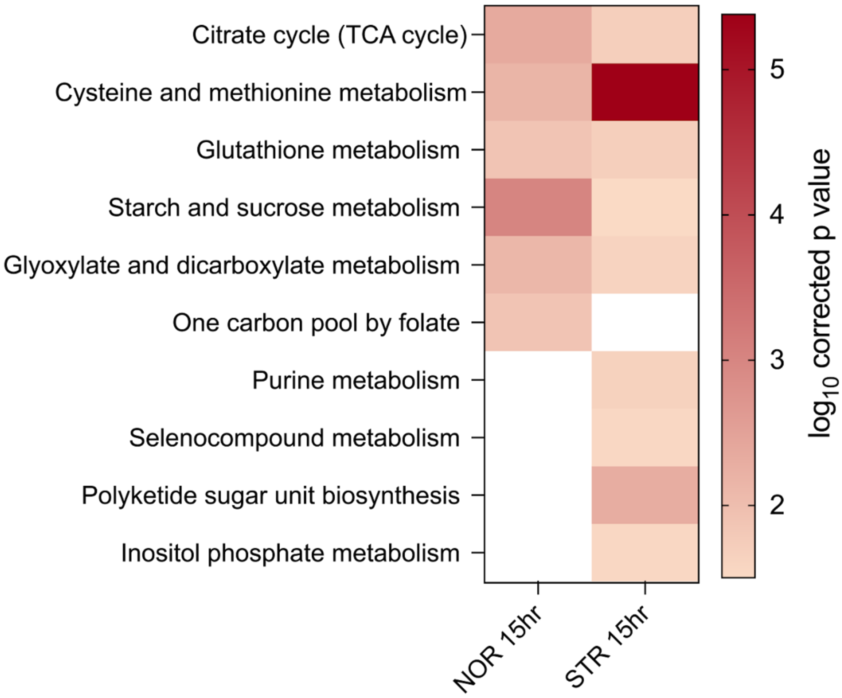


**S3.** The scatter plot (A) depicts the Pearson correlation analysis of proteins commonly identified upon norfloxacin and streptomycin treatment. Each dot represents a single protein with its $\log_2$ fold change for NOR (x-axis) and STR (y-axis) treatment at t = 25 hour time point. Heatmap (B) depicts the KEGG pathway enrichment analysis of the proteins upregulated and exclusively expressed in response to NOR and STR at 15 hours.

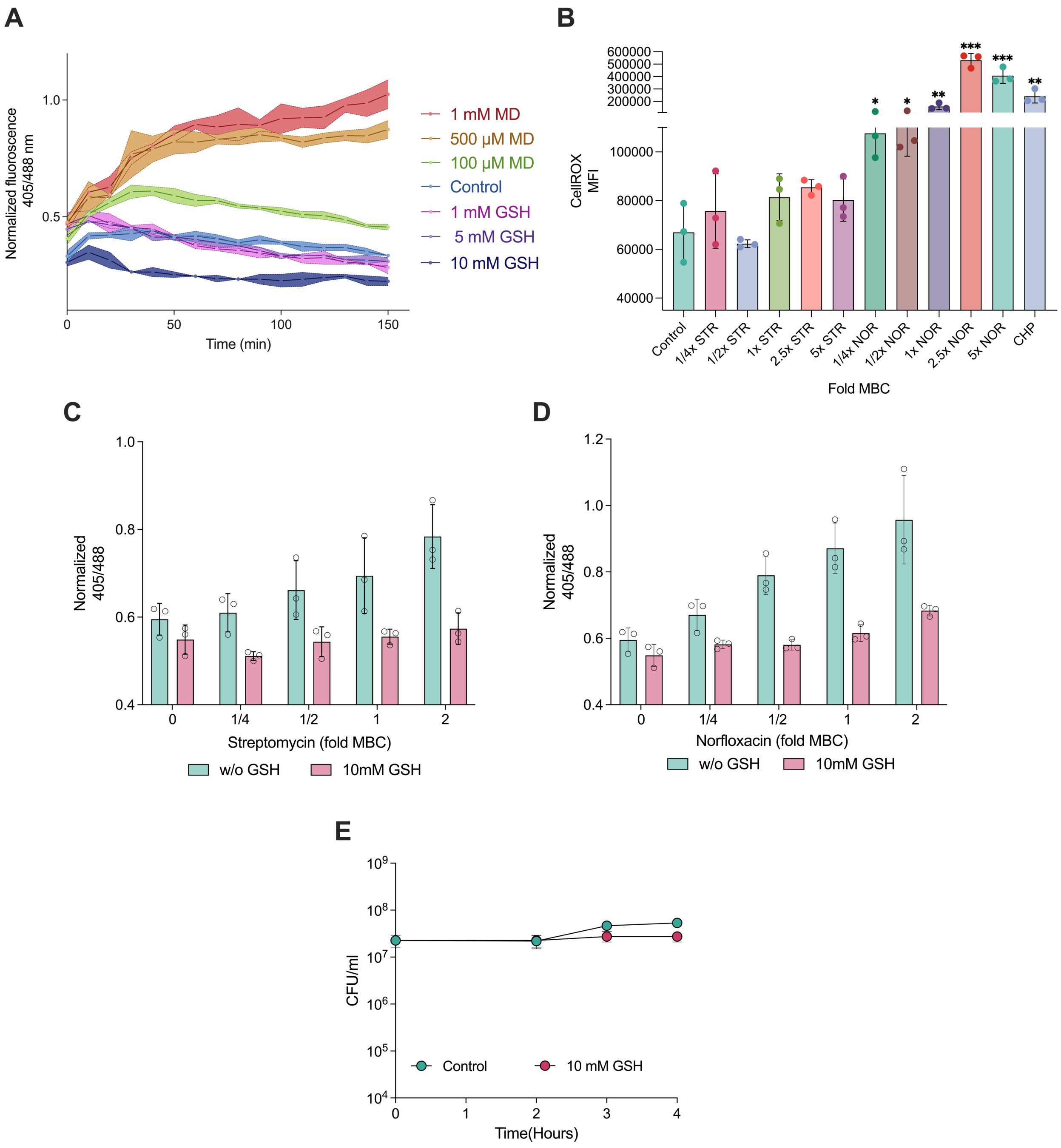


**S4.** (A) Time-course ratio metric response of *M. smegmatis* expressing Mrx1-roGFP2 redox biosensor, at increasing concentrations of menadione (MD) and glutathione (GSH). Mean fluorescence intensities of CellROX deep red dye (B) upon 3 hours of treatment with varying concentrations of norfloxacin and streptomycin; 5mM cumene hydroperoxide (CHP) was used as a positive control. Ratiometric response of *M. smegmatis* in response to increasing concentrations of streptomycin (C) and norfloxacin (D) for 3 hours, with and without co-treatment with 10 mM glutathione. (E) Effects of GSH alone on the survival of *M. smegmatis*. *$p < 0.05$, **$p < 0.01$, ***$p < 0.001$ were calculated by student's t-test (unpaired).

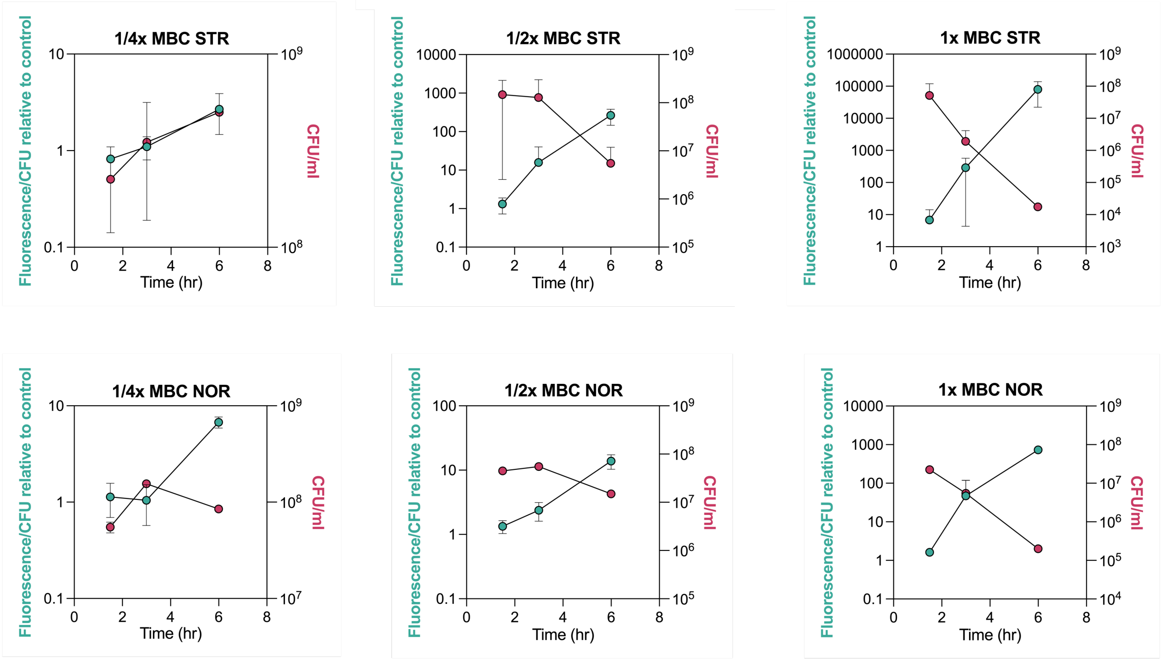


**S5.** Figures represent the resazurin fluorescence (at 530 nm excitation and 590 nm emission) on left y-axis and viability of cells in right y-axis, in *M. smegmatis* exposed to either ¼x, ½x, or 1x-MBC$_{99}$ of STR and NOR, respectively.

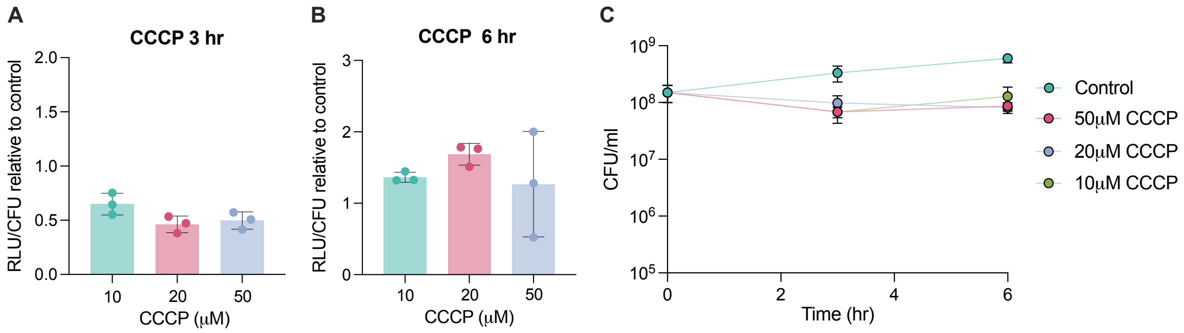


**S6.** Bar graph depict the effect of CCCP treatment alone on relative luminescence (RLU) measurements indicating intracellular ATP levels (A and B), and mycobacterial viability (C), at indicated time points. Data represent the mean of three biological replicates.

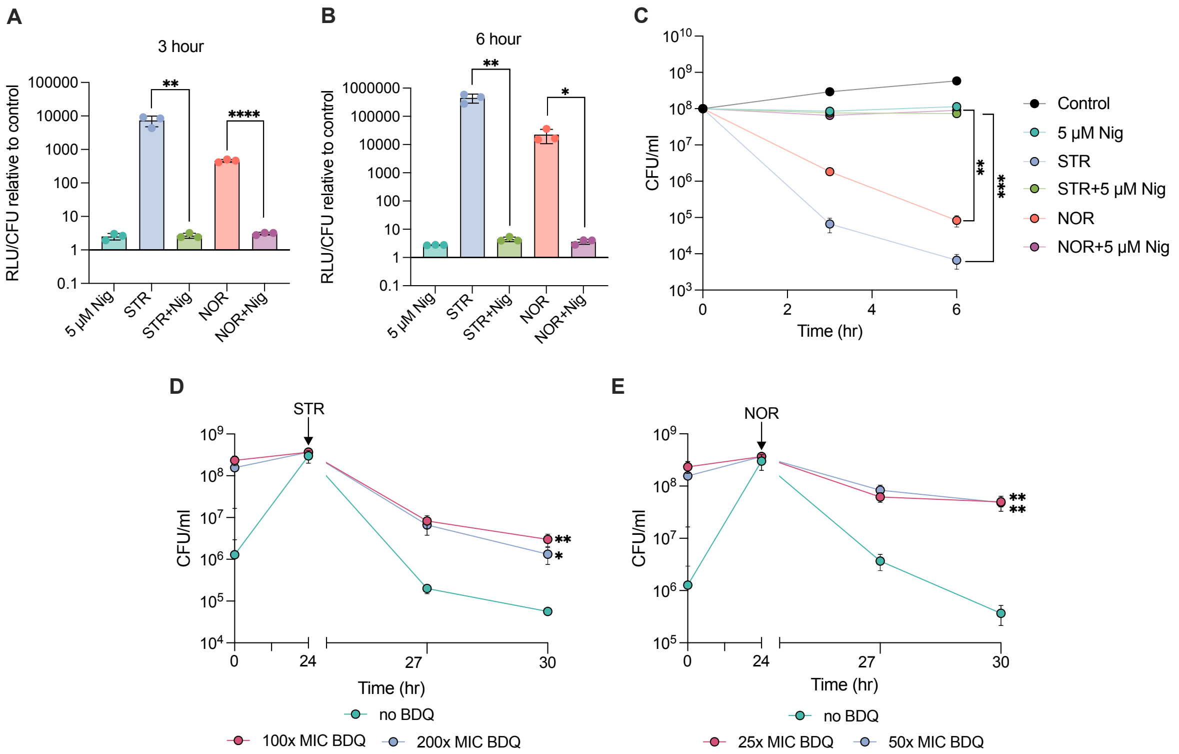


**S7.** Bar graphs depict relative luminescence (RLU) measurements indicating intracellular ATP levels (A and B) in response to 5 µM nigericin (Nig) alone and in co-treatment with 1x $MBC_{99}$ NOR or STR for indicated time points. The effect of nigericin co-treatment on the survival of *M. smegmatis* for up to 6 hours (C). Figures (D and E) depict time-kill curve of *M. smegmatis* pre-treated with BDQ for 24 hours, and subsequently challenged with STR and NOR, respectively. 1x $MIC_{99}$ of BDQ used in the study was 0.0075 µg/ml. All data points represent the mean of at least three independent replicates ± SD. Statistical significance was calculated by students' t-test (unpaired), $^*p < 0.05$, $^{**}p < 0.01$, $^{***}p < 0.001$, $^{****}p < 0$.

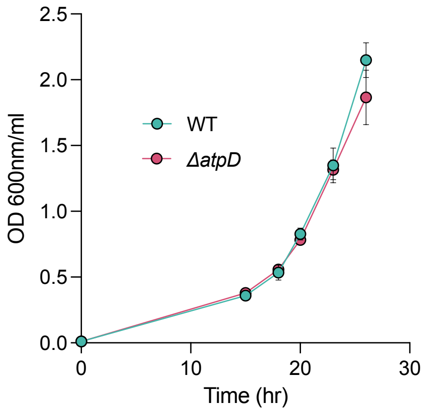


**S8.** Figure depicts the growth curve of wild type and Δ*atpD* strain of *M. smegmatis*. Data represent the mean of three biological replicates.

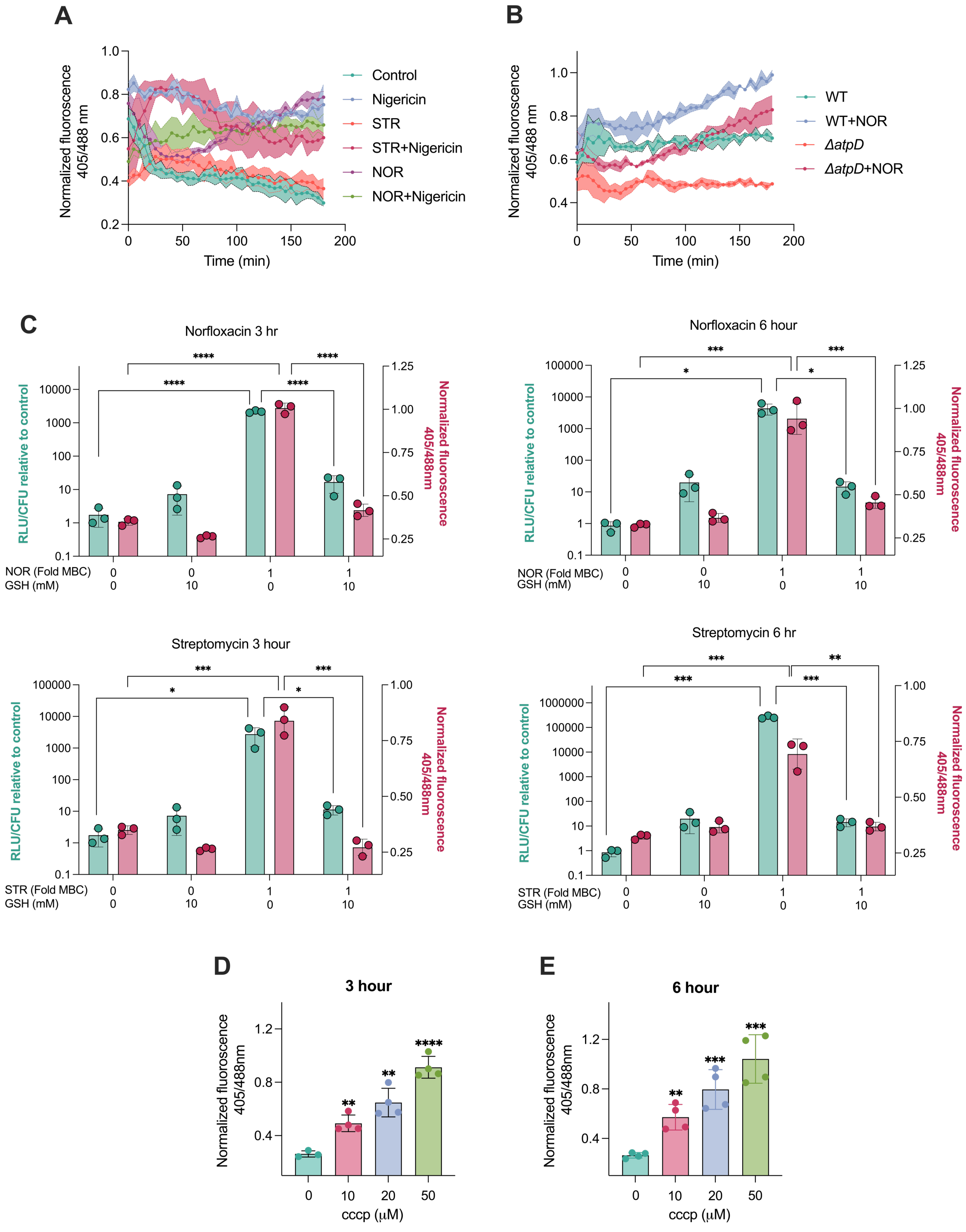


**S9.** Line graph (A) represent the ratio metric response of *M. smegmatis* expressing Mrx1-roGFP2 redox biosensor for 3 hours in response to 5 µM nigericin alone and in co-treatment with 1x $MBC_{99}$ NOR or STR. Figure (B) represent the ratio metric response of wild type and Δ*atpD* ($Hyg^R$) strain of *M. smegmatis* expressing Mrx1-roGFP2 redox biosensor ($kan^R$) in the presence and absence of 1x $MBC_{99}$ NOR. Bar graph (C) illustrates the effect of co-treatment of 10 mM GSH on antibiotic-induced ATP levels - the figure depicts the relative luminescence

(RLU) measurements indicating intracellular ATP as well as ratio metric response of *M. smegmatis* expressing Mrx1-roGFP2 redox biosensor in response to 1x $MBC_{99}$ of NOR or STR alone, 10 mM GSH alone or in combination, after 3 and 6 hours of treatments. Bar graphs depict the ratio metric response of *M. smegmatis* expressing Mrx1-roGFP2 redox biosensor in response to increasing concentrations of CCCP alone after 3 hours (D) and 6 hours (E). All data points represent the mean of at least three independent replicates ± SD. Statistical significance was calculated by students' t-test (unpaired), *$p < 0.05$, **$p < 0.01$, ***$p < 0.001$, ****$p < 0$.

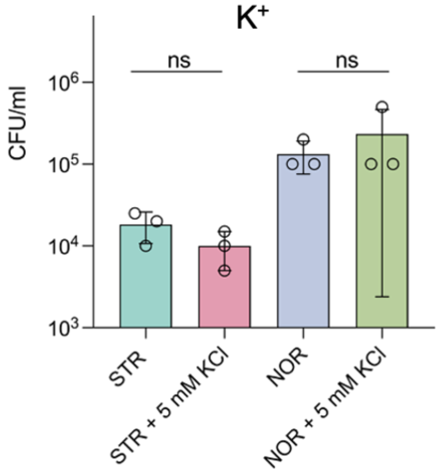


**S10**. Effect of 5 mM potassium chloride on the survival of *M. smegmatis* challenged with 1X $MBC_{99}$ of STR and NOR for 6 hours. Co-treatment with monovalent metal ions (which is not sequestered by ATP) do not provide rescue against STR and NOR lethality.

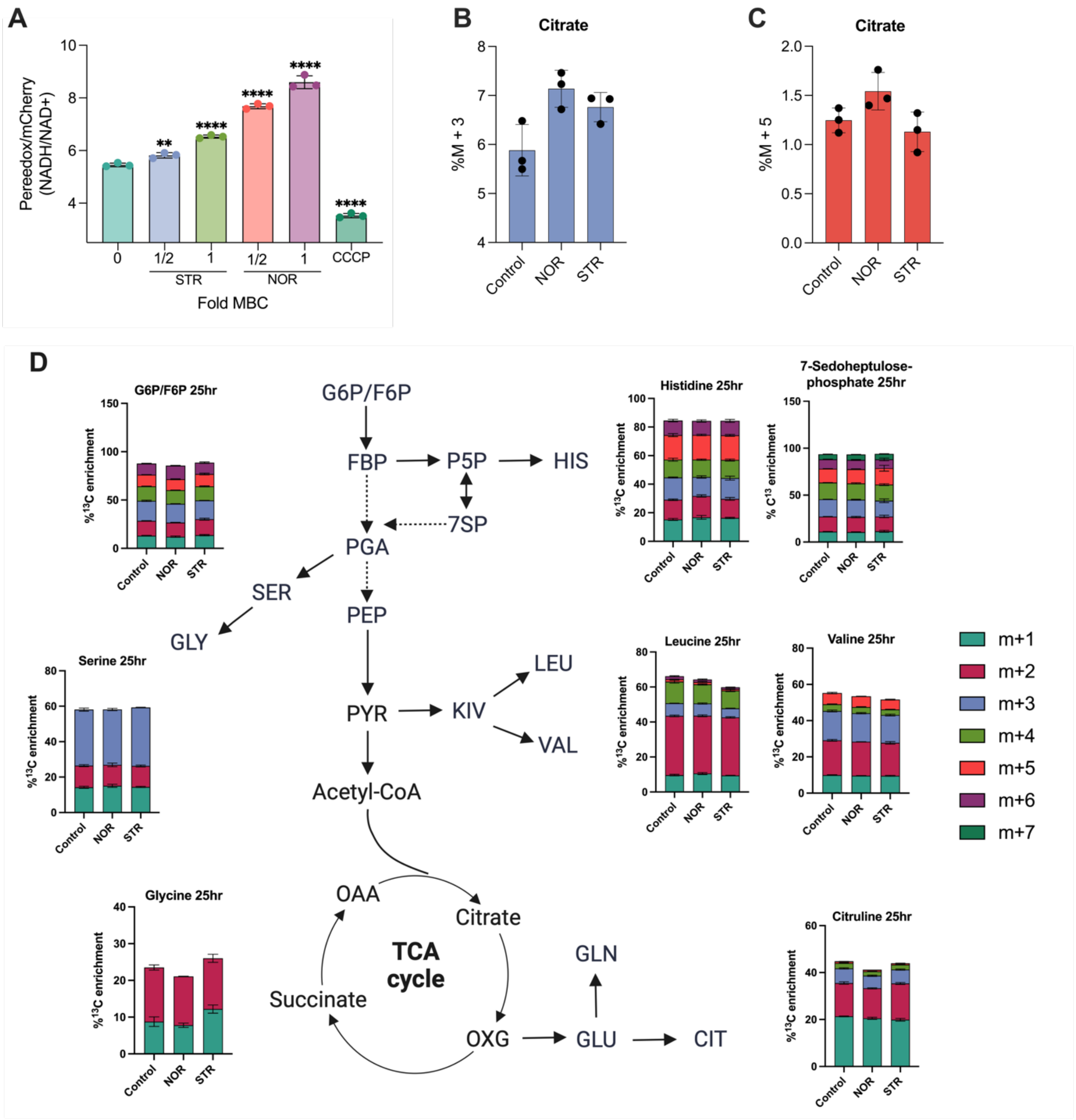


**S11.** Bar graph (A) depicts the Peredox/mCherry ratio (NADH/NAD+) of the NADH biosensor in *M. smegmatis* exposed to increasing concentrations of STR and NOR; 50 micromolar CCCP was used as a control. Figures depict the % $^{13}$C enrichment of M+3 (B) and M+5 (C) species of citrate. Figure (D) illustrates the % $^{13}$C enrichment in metabolites synthesized from central carbon metabolism of *M. smegmatis* challenged with ¼ xMBC$_{99}$ of NOR and STR at 25 hour time point (recovery phase). All data represent the mean ± SD from independent triplicates. *p < 0.05, **p < 0.01, ***p < 0.001 were calculated by student's t-test (unpaired).
